# Mental simulations and action language are impaired in individuals with aphantasia

**DOI:** 10.1101/2023.03.16.532945

**Authors:** W. Dupont, C. Papaxanthis, F. Lebon, C. Madden-Lombardi

**Author notes:** Corresponding author : William Dupont. Equal contribution (F. Lebon, C. Madden-Lombardi).

## Abstract

Action reading is thought to engage motor simulations, yielding modulations in activity of motor-related cortical regions, and contributing to action language comprehension. To test these ideas, we measured 1) corticospinal excitability during action reading, and 2) reading comprehension ability, in individuals with normal and impaired imagery (i.e., phantasia and aphantasia, respectively). Thirty-four participants (17 phantasic and 17 aphantasic) were asked to read manual action sentences. By means of transcranial magnetic stimulation, we triggered motor-evoked potentials (MEP) in the target right index finger. MEP amplitude, a marker of corticospinal excitability, increased during action reading relative to rest for phantasic individuals, but not for aphantasic individuals. This result provides neurophysiological evidence that individuals living with aphantasia present a real neurophysiological deficit in motor system engagement during action reading. Furthermore, deep-level reading comprehension ability was impaired in individuals with aphantasia, who had difficulty selecting words that best fit the context of sentences. Altogether, these findings support the idea that motor simulations, along with the activation within the motor system, contribute to action language comprehension.

## Introduction

The generation of mental simulations is a fundamental characteristic of human existence, allowing us to visualize objects, remember past events or predict the sensorimotor consequences of an action. A growing body of evidence suggests that understanding action sentences exploits the neural mechanisms of mental simulation used for perception and action. According to the embodied view of language comprehension (Barsalou, 2008; Fischer and Zwaan, 2008; Gallese and Lakoff, 2005; Zwaan and Madden, 2005), action language would automatically and unconsciously elicit a motor simulation during action reading, in order to capture features of described action, such as direction and duration (Glenberg and Kaschak, 2002; Kaschak et al., 2005; Matlock, 2004), and the target effector (Bergen et al., 2004, 2003). These motor simulations may serve to more efficiently understand the action described in the sentences (Bailey, 1998; Barsalou, 1999).

Neurophysiological manifestations of such simulation during action reading include the increase of corticospinal excitability in Transcranial Magnetic Stimulation (TMS) studies (W. Dupont et al., 2022a; Innocenti et al., 2014; Labruna et al., 2011; Papeo et al., 2013, 2009) and the activation of the motor system in neuro-imaging studies (Aziz-Zadeh et al., 2006; Hauk et al., 2004; Tettamanti et al., 2005; Van Dam et al., 2010; Wu et al., 2013). Although certain findings speak against the automatic involvement of the motor cortex during the processing of action words (de Zubicaray et al., 2013; Grossman et al., 2002; Longe et al., 2007; Mahon, 2015; Mahon and Caramazza, 2008; Perani et al., 1999; Wurm and Caramazza, 2019), these discrepancies could be partially explained by task-related factors, such as the type of linguistic tasks or stimuli used, or by top-down processes, such as participants’ reading strategies. Tomasino et al. (2008) suggested that the involvement of the primary motor cortex (M1) during action reading may depend on the (explicit or implicit) strategy employed by individuals to generate a motor simulation of the action described in the sentence (Tomasino and Rumiati, 2013). This is consistent with Ferreira and colleagues’ claim that representations activated during language processing “are only good enough to tackle the task at hand and become elaborated only if mandated by the task at hand” - what they call “good enough” representations (Ferreira et al., 2002). Thus, depending on the motivation and goals of the reader as well as the task constraints (time limits, sematic depth, etc.), motor simulations may be activated to a greater or lesser degree.

Current neurophysiological and behavioral evidence remains insufficient to support the idea that motor simulations associated with the activation of the motor system contribute to the comprehension of described actions. Individuals with an inability to explicitly generate mental images would provide an ideal testbed to measure whether the motor system is recruited during action reading and whether it plays a role in language comprehension. The present paper aims to shed new light on this issue by exploring the influence of motor simulations during action reading in individuals living with an aphantasia. The term “aphantasia” refers to an actual inability to voluntarily imagine an event or a movement (Zeman et al., 2015). Francis Galton (1880) first reported that some individuals have a weaker ability or even an inability to imagine. This idea that some individuals are unable to create visual images in their mind, without any associated disorders, has recently reemerged and seems to be attracting attention. Although researchers have also reported physiologic evidence of a real imagination inability, using binocular rivalry and eye tracking methodologies (Kay et al., 2022; Keogh and Pearson, 2018), this phenomenon is mainly evaluated by subjective reports of visual imagery vividness (Dawes et al., 2020; Milton et al., 2021; Zeman et al., 2020, 2016, 2015, 2010).

In the current study, we probed the ability of phantasic and aphantasic individuals to create mental images of actions. We collected subjective reports of imagery vividness and we measured corticospinal excitability by means of TMS during action reading. To determine whether the inability to create mental images while reading impacted comprehension ability, we tested the memorization of text elements (surface, or memory-based comprehension) and the inferences constructed by the reader (deep, or context-based comprehension). If a motor simulation is accompanied by the activation of the motor system during action reading, and if this simulation contributes to reading comprehension, we expect to observe a lack of corticospinal increase during action reading for aphantasics, accompanied by reduced reading comprehension ability.

## Results and Discussion

### Explicit imagery ability questionnaires

We confirmed that aphantasics (n=16) reported having difficulties or even the inability to explicitly create mental images of common actions, in comparison to phantasics (n=16). Average scores on the Vividness of Movement Imagery Questionnaire-2 (VMIQ-2, Roberts et al., 2008) and at the Spontaneous Use of Imagery Scale (SUIS, Ceschi and Pictet, 2018) approached the boundary of no imagery for aphantasics (see Figure 1 for main results and Table 1 in supplementary section for details). These subjective reports are in line with the literature (Dawes et al., 2020; Keogh and Pearson, 2018; Zeman et al., 2015).

**Table 1:** Average scores on Vividness of Movement Imagery Questionnaire-2 (VMIQ-2, Roberts et al., 2008) and Spontaneous Use of Imagery Scale (SUIS, Ceschi and Pictet, 2018) for aphantasic and phantasic participants. The worst and best score for each modality on the VMIQ-2 is 60 and 12, respectively. The worst and best score for the SUIS questionnaire is 12 and 60, respectively. A repeated measures ANOVA on VMIQ-2 scores pointed out a main effect of Group (F_1,30_=385.64, p<0.0001, ηp^2^=0.927), with a greater imagery ability for phantasic (24.64 ±8.58; Cohen’s d=5.12) than aphantasic participants (57.90 ±5.52). We observed a Group x Perspective interaction (F_2,60_=4.606, p=0.013, ηp^2^=0.133), demonstrating a marginally significant difference between the internal (20.68 ±5.81) and kinaesthetic (26.75 ±8.33, p=0.080, Cohen’s d=0.87) vividness of visual motor imagery in phantasics, and no such difference in aphantasics. Moreover, an independent T-test on SUIS scores revealed a greater utilization of imagery in everyday life for phantasic individuals (40.50 ±7.03) than that for aphantasic participants (18.62 ±4.57; t(_30_)=10.42, p<0.0001, Cohen’s d=3.81)

| Questionnaires | VMIQ-2 |  |  | SUIS |
| --- | --- | --- | --- | --- |
|  | External visual | Internal visual | Kinaesthetic |  |
| **Phantasic individuals (Mean ±SD)** | 26.50<br>±<br>10.11 | 20.68<br>±<br>5.81 | 26.75<br>±<br>8.33 | 40.50<br>±<br>7.03 |
| **Aphantasic individuals (Mean ±SD)** | 59.00<br>±<br>3.09 | 59.00<br>±<br>1.89 | 55.68<br>±<br>8.62 | 18.62<br>±<br>4.57 |

### Implicit imagery during action reading

To probe whether implicit imagery is at play during action reading, participants were asked to self-report the vividness of images while reading about concrete visual, concrete motor, and abstract content on a scale of 1 to 5, 1 being a highly vivid image and 5 being no image (Madden-Lombardi et al., 2015). A repeated measures ANOVA pointed out a main effect of Group (F_1,30_=326.60, p<0.001, ηp^2^=0.915), and of Content (F_2,60_=69.268, p<0.001, ηp^2^=0.697) and an interaction between Group and Content (F_2,60_=60.154, p<0.001, ηp^2^=0.667; Figure 2). Phantasics reported easily picturing concrete motor sentences (average score: 1.47 ±0.43) and visual sentences (1.70 ±0.49) during reading. As expected, their scores for abstract sentences (3.57 ±0.77) differed from concrete motor and visual sentences (p<0.001 and Cohen’s d=3.48; p<0.001 and Cohen’s d=2.99, respectively). On the contrary, aphantasics reported very little vividness for all items during reading (abstract sentences: 4.77 ±0.53; concrete motor sentences: 4.70 ±0.37; visual sentences: 4.69 ±0.33), indicating that they did not picture the events while reading. Their scores were significantly different from those of phantasics (abstract sentences: Cohen’s d=1.88; p<0.001 concrete motor sentences: Cohen’s d=8.42, p<0.001; and visual sentences: Cohen’s d=7.39, p<0.001).

**Figure 2:**
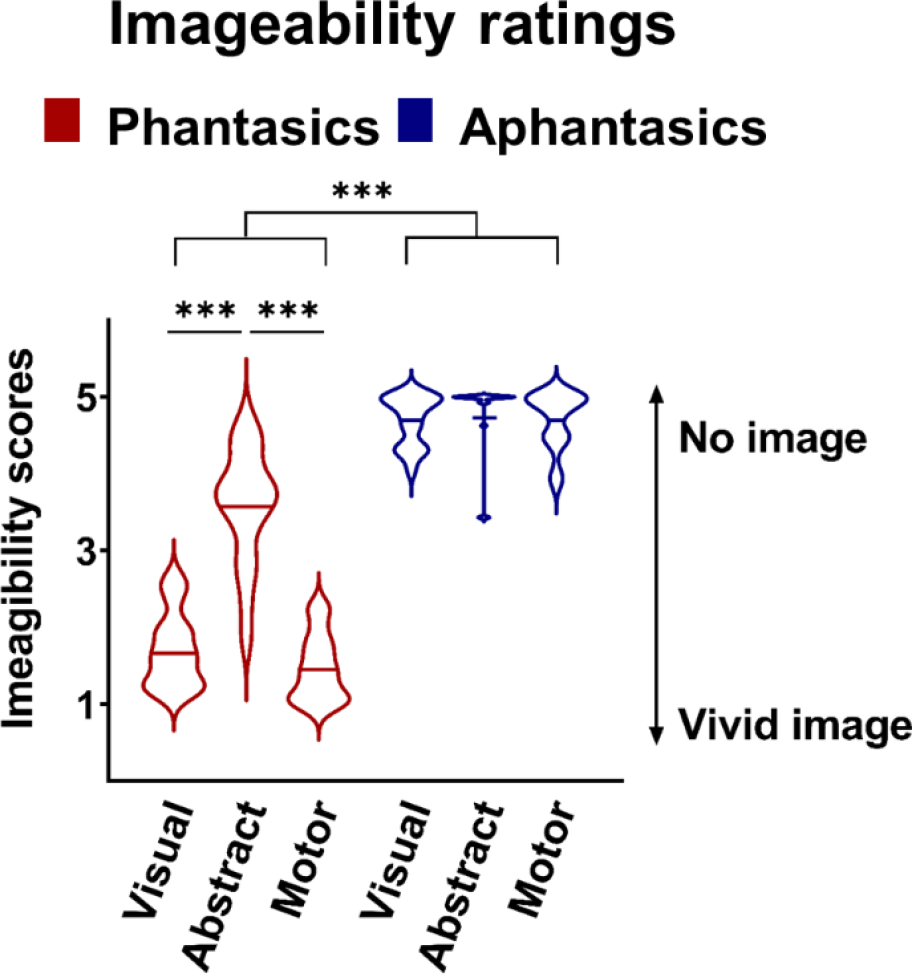
Violin plots of imageability ratings of visual, abstract and motor sentences for phantasics and aphantasics. Thick horizontal lines indicate means. *** = p<0.001.

Next, we measured corticospinal excitability while participants read action sentences involving the hand. As expected, MEP amplitude increased during reading in comparison to rest for phantasics (29.51 ±24.81%, t=4.758, p<0.001), but not for aphantasics (3.07 ±14.35%, t=0.857, p=0.404). The percentage of MEP amplitude increase was statistically different between the 2 groups (t(_30_)=3.689, p=0.001; Figure 3). This result extends previous research, suggesting a clear link between the activation of the primary motor cortex and the motor simulation while reading about actions (Barsalou, 2008; Fischer and Zwaan, 2008; Gallese and Lakoff, 2005; Jeannerod, 2008).

**Figure 4:**
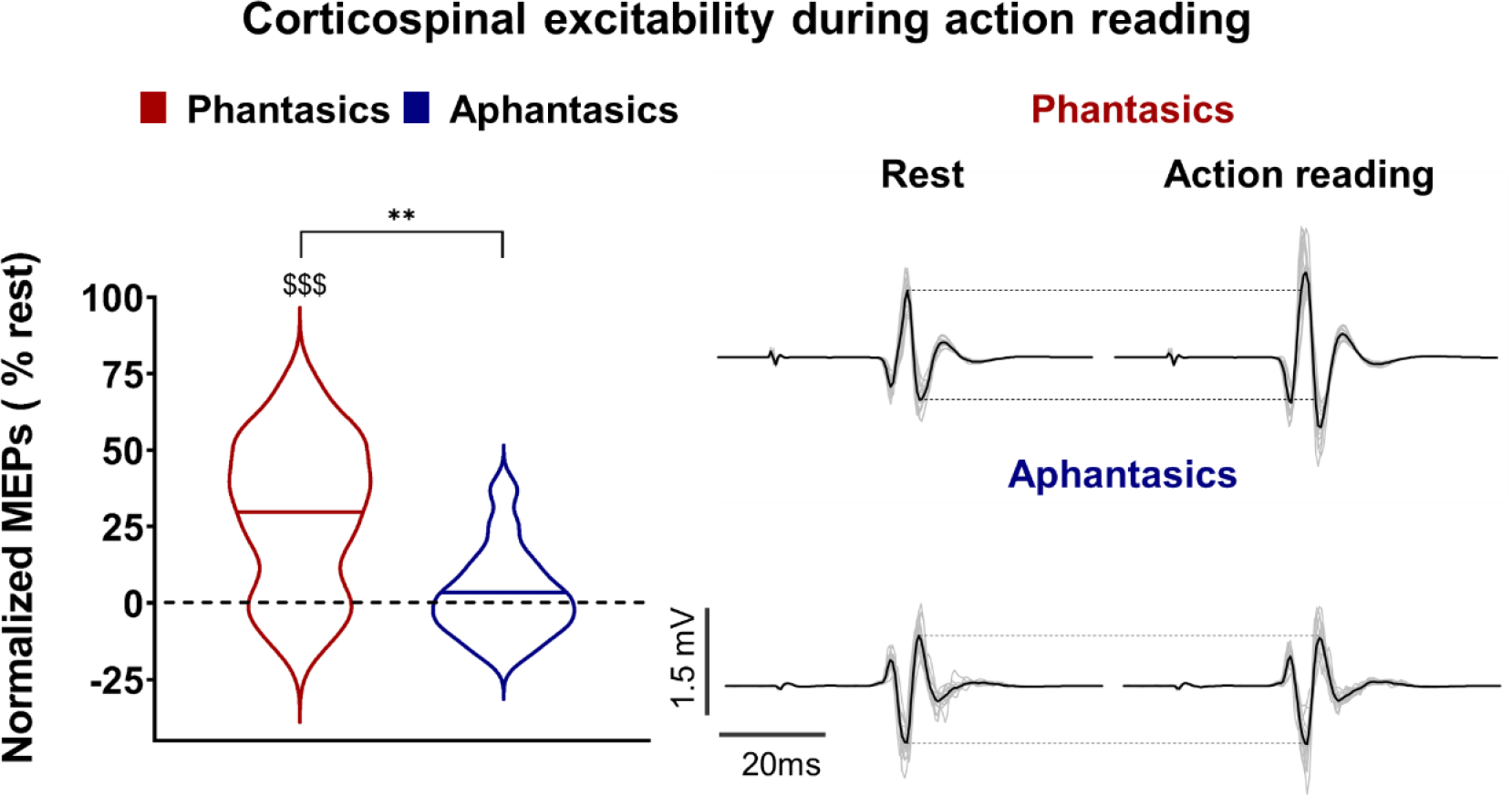
Corticospinal excitability during action reading for phantasics and aphantasics. Violin plots on the left side represent MEPs normalized to rest during action reading. Thick horizontal lines mark the mean. The right side of the panel illustrates raw MEPs of a typical subject (grey lines). The black line is the average MEP of the condition for this participant. **=p<0.01 indicates a significant difference between the two groups and $$$=p<0.01 indicates a significant difference from zero (rest).

### Reading comprehension

Finally, we investigated whether aphantasics’ inability to mentally simulate altered their reading comprehension. We assessed low and high level reading comprehension abilities, characterized respectively by the memorization of text elements (memory-based comprehension) and the inferences constructed by the reader (context-based comprehension). We analyzed the percentage of correct responses. Whereas memory-based reading comprehension scores did not differ between phantasics (80.63 ±12.43%) and aphantasics (73.75 ±12.04%; t(_30_)=1.58, p=0.12, Cohen’s d=0.58), context-based comprehension scores were higher for phantasics (87.69 ±6.14%) than for aphantasics (79.63 ±9.54%; t(_30_)=2.84, p=0.008, Cohen’s d=1.04). This later result complements previous research reporting that the vividness of mental simulations can influence reading comprehension (Denis, 1982). Taken together, these findings suggest that the inability to simulate a described action may impair reading comprehension.

### General discussion

The present study provides several notable findings related to motor simulation and action reading. First, our self-report data replicates and extends previous research demonstrating a deficit in voluntary mental simulation for aphantasics. Second, the lack of increase in corticospinal excitability during the passive reading of action phrases in aphantasics demonstrates that these individuals present a real neurocognitive impairment, not under the influence of their own strategies, volition, or metacognition. Finally, aphantasics’ impaired performance on the context-based reading comprehension task sheds light on the role of mental simulation in language comprehension.

Concerning the first point, while previous research has demonstrated that aphantasics are less able (or unable) to engage in various types of mental simulation (Dawes et al., 2020; Keogh and Pearson, 2018; Zeman et al., 2015), our self-report estimates of vividness on the VMIQ-2 extend these results to the realm of motor simulation. Furthermore, our imageability ratings are, to our knowledge, the first to illustrate that aphantasia also encompasses an inability to automatically elicit motor or visual simulations during the passive reading of action sentences. This result extends recent investigations demonstrating the deterioration of implicit simulations during mental rotation (Milton et al., 2021; Pounder et al., 2022; Zeman et al., 2010; Zhao et al., 2022), involuntary simulations during night-time dreams (Dawes et al., 2020; Zeman et al., 2015, 2010; see review: Whiteley, 2020), or action observation (Dupont et al., preprint 2022c).

Importantly, we reinforced these self-report data from questionnaires with neurophysiological measures. We observed that, relative to rest, action reading increased corticospinal excitability in phantasics but not aphantasics (for similar resuls in phantasic participants, see Dupont, 2022; Innocenti et al., 2014; Labruna et al., 2011; Papeo et al., 2013, 2009). Indeed, in the case of aphantasia, the reported absence of a motor simulation is coupled with a real neurological deficit, manifest in the lack of an increase in corticospinal excitability during action reading. These findings suggest that the activation of the motor system during action reading is directly related to the simulation of the described action. This is consistent with the idea that motor system activation is not guaranteed upon reading or hearing action language, but rather depends on whether subjects engage in motor imagery (automatically or with effort) during the reading task (Ferreira et al., 2002; Tomasino and Rumiati, 2013).

Finally, the results of the current study suggest that the inability to mentally simulate is accompanied by a deficit in reading comprehension in aphantasics compared to phantasics. This is consistent with previous research showing that voluntarily engaging mental simulations elicited during reading can improve comprehension and contribute to representations that remain available for subsequent cognitive tasks (Chaguiboff and Denis, 1981; Commodari et al., 2020; Denis, 1982; Guarnera et al., 2019; Kulhavy and Swenson, 1975; Pressley, 1976). We propose that distributed activation in sensory-motor areas contributes richness to the comprehended meaning, adding to the lexical access and associative meaning afforded by treatment in typical language areas. Therefore, it is not surprising that scores on surface-level or memory-based comprehension would be spared while the deeper contextual comprehension is impaired in the case of aphantasia. The motor simulation would add richness to better guide contextually appropriate choices on the context-based comprehension task, but not necessarily increase chances of answering correctly on a recognition test. This might seem to contradict previous studies reporting that mental simulation plays a significant role in memorizing a text (Cayol et al., 2020; Chaguiboff and Denis, 1981; Denis, 1982), and that aphantasic individuals exhibit (especially episodic) memory impairments (Bainbridge et al., 2021; Dawes et al., 2020; Milton et al., 2021; Zeman et al., 2020). However, a lack of simulation would likely affect recall memory that is based on a deeper level of comprehension rather than mere word recognition. Further research is underway to better understand the extent of impairments caused by the inability to simulate.

To conclude, our results provide novel neurophysiological evidence that aphantasics’ subjectively reported deficit in mental simulation is accompanied by a lack of activation in the motor system, as well as a measurable impairment in reading comprehension behavior. Second, our results suggest that action reading does not automatically engage the primary motor cortex in all cases. Implicit or explicit generation of motor imagery seems to be necessary to activate the motor system during action reading, and there exists a certain segment of the population in which this does not occur.

## Material and methods

### Participants

Thirty-four right-handed aphantasic (n=17) and phantasic (n=17) participants were included in the study. Two participants were excluded from the analysis (Data and statistical analysis). We performed the statistical analyses with 16 aphantasics (9 women, mean age: 20; range: 18-26) and 16 phantasics (6 women, mean age: 23; range: 19-26). We ensured the right laterality with the Edinburgh inventory (Oldfield, 1971). All participants were French native speakers and completed the questionnaire by Lefaucheur et al., 2011 prior to participation to determine whether they were eligible for TMS. An Ethics Committee (CPP SOOM III, ClinicalTrials.gov Identifier: NCT03334526) in accordance with the Declaration of Helsinki approved all procedures (excluding pre-registration).

### Procedure and stimuli

The participants came to the laboratory for two experimental sessions. The first behavioral session included subjective assessments of mental simulation abilities, as well as reading comprehension tests. Then, individuals followed a neurophysiological session with TMS delivered at rest (fixation cross) and during action reading (200, 300, 400, 500 ms). Participants sat in an armchair and read stimuli presented on a 19-inch LCD monitor by a home-made software (Neurostim) which controlled TMS triggering and synchronized electromyographic recordings.

#### Imagery questionnaires

All participants completed the Vividness of Movement Imagery Questionnaire-2 (VMIQ-2) (Roberts et al., 2008), the Spontaneous Use of Imagery Scale (SUIS) (Ceschi and Pictet, 2018), and an imageability ratings assessment during reading (Madden-Lombardi et al., 2015). For the VMIQ-2, the participants imagined multiple actions/movements using three modalities (External Visual Imagery, Internal Visual Imagery, Kinaesthetic imagery), and then rated how vivid their imagery was for each movement on a scale of 1-5 (with 1 = “Perfectly clear and vivid as normal vision” and 5= “No image at all, you only think about the movement”). For the SUIS, we evaluated the general tendency of individuals to use visual mental imagery in everyday situations. Each item is rated on a 5-point scale (from 1 = “never appropriate”; to 5 = “always completely appropriate”) according to the item described and the individual’s functioning. The reading imageability ratings assessment was adapted from the revised version of the Vividness of Visual Imagery Questionnaire (VVIQ-2; (Madden-Lombardi et al., 2015; Marks, 1995). Here, participants read sentences evoking abstract, visual or motor contents and were asked to rate how vivid a representation arose in their minds while reading (from 1 = “highly vivid image”; to 5: “nonvivid image”).

#### Reading comprehension assessments

Reading comprehension assessments may focus on the memorization of text features and components, characterizing the “low-level” or “surface level” comprehension. Alternatively, assessments may focus on the inferences constructed by the reader, which are related to a “deep” or “high-level” of comprehension. To investigate memory-based comprehension, participants performed a “Remember versus Know” paradigm (Tulving, 1985), in which they had to identify whether words were encountered in a previously read text (yes or no response), and in what context they encountered that word. In this task, we measured on 80 trials the percentage of correct responses. To assess higher context-level comprehension, we used a Curriculum-Based Measurement-Maze (Parker et al., 1992), in which readers relied on inferences to select a contextually-appropriate word from three alternatives. After the first sentence, every seventh word in the passage was replaced with that correct word and two distracters. While distractors did not violate any sentential (grammatical or semantic) rules, participants had to use the context to select the word that best fit with the rest of the passage. Participants made 31 choices to complete the 10 sentences, and we measured the percentage of correct responses.

Then, to ensure that our sample of aphantasics and phantasics had similar cognitive abilities, we measured performance on mental arithmetic and Raven’s matrix tasks (%). For the Raven matrix task, participants were asked to complete the missing part of a figure with one of the four propositions. For the mental arithmetic task, participants were asked to complete calculations without any support. Each task contained 14 trials.

#### Action reading stimuli (transcranial magnetic stimulation session)

Sixty-four French sentences were generated that referred to hand actions (e.g., “I have a hair on my arm, I pull it out”; “J’ai un cheveu sur mon bras, je le retire”). All sentences were presented in the first-person present tense and were created so that the target verb occurred at the end of the sentence. This final pronoun-verb segment was presented on a subsequent screen after the sentence beginning was presented (Figure 5). All stimuli order was randomized and counterbalanced. We randomly delivered TMS pulses at 200 ms, 300 ms, 400 ms, and 500 ms after the appearance of the action verb (16 TMS pulses per condition). Moreover, sixteen TMS pulses were delivered at rest (fixation cross), which served as reference stimulations.

Surface Electromyography (EMG) was recorded using 10-mm-diameter surface electrodes (Contrôle Graphique Médical, Brice Comte-Robert, France) placed over the right First Dorsal Interosseous (FDI) muscle. Before positioning the electrodes, the skin was shaved and cleaned to reduce EMG signal noise (<20μV). EMG was amplified with a bandpass filter (10-1000 Hz) and digitized at 2000Hz (AcqKnowledge; Biopac Systems, Inc., Goleta, CA). We calculated the root mean square EMG signal (EMGrms) for further analysis.

Using a figure-of-eight coil (70 mm diameter) connected to Magstim 200 stimulator (Magstim Company Ltd, Whitland), single-pulse TMS were delivered over the motor area of the right FDI muscle. The coil rested tangentially to the scalp with the handle pointing backward and laterally at 45° angle from the midline. Using a neuronavigation system (BrainSight, Rogue Research Inc.) with a probabilistic approach (Sparing et al., 2008), the muscle hotspot was identified as the position where stimulation evoked the highest and most consistent MEP amplitude for the FDI muscle. This position was determined via a regular grid of 4 by 4 coil positions with a spacing of 1 cm (centered above the FDI cortical area x=-37, y=-19, z=63; Bungert et al., 2017; Sondergaard et al., 2021). During the experiment, the intensity of TMS pulses was set at 130% of the resting motor threshold, which is the minimal TMS intensity required to evoke MEPs of 50µV peak-to-peak amplitude in the right FDI muscle for 5 trials out of 10 (Rossini et al., 2015).

**Figure 4:**
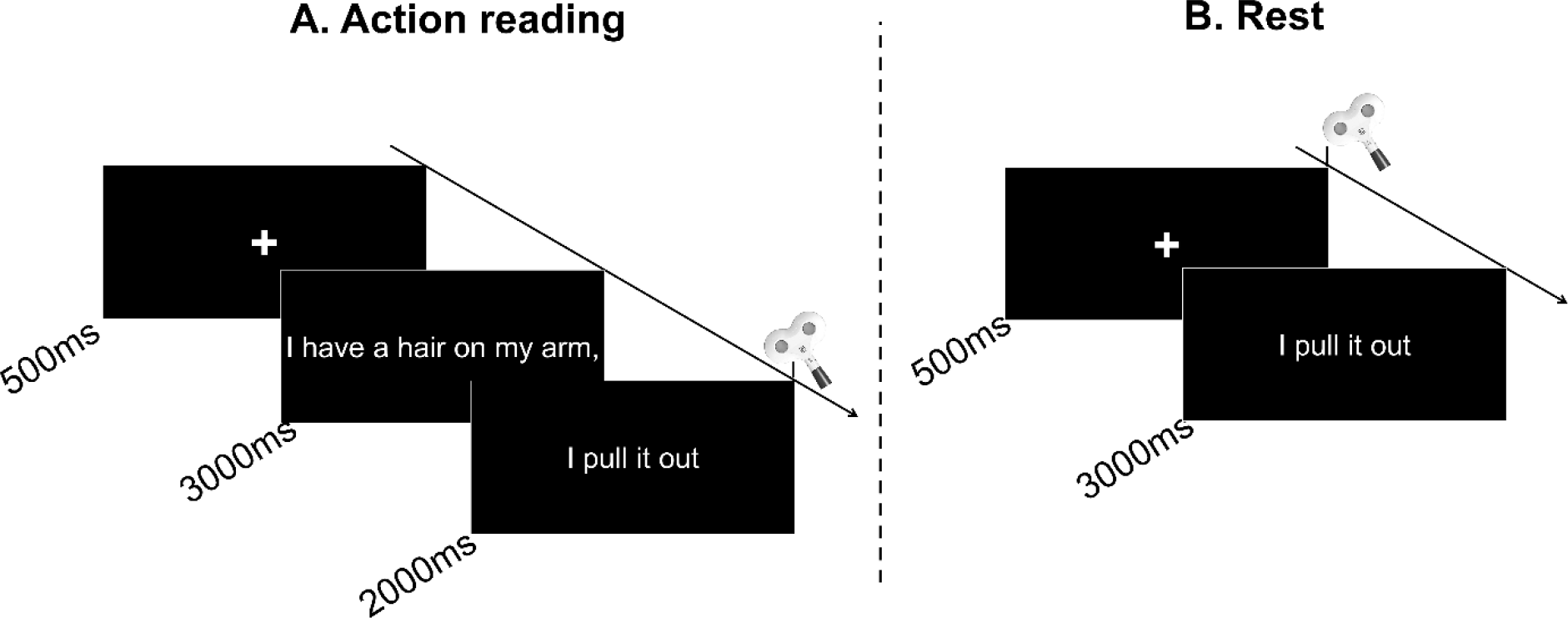
Experimental procedure. Each trial started with a fixation cross to indicate the beginning of a trial. **A. Action reading**. The participants silently read sentences and we randomly delivered TMS pulses at 200 ms, 300 ms, 400 ms, and 500 ms after the appearance of the action verb (third screen). **B. Rest**. The participants silently read stimuli and we delivered TMS pulses at the fixation cross.

### Data and statistical analysis

First, based on a large effect size between the action sentences and the rest condition from a previous study (Dupont et al., preprint 2022b), we estimated via G* Power (version 3.1.9.4., Faul et al., 2007), that 16 participants per group would be needed for analysis. Specifically, we used a Cohen’s d of 0.589 from a t-test (action reading compared to rest), and a power of 0.95 to find 16 participants per group for a 2×2 interaction (group x task) with an ANOVA.

Matlab (The MathWorks, Natick, Massachusetts, USA) was employed to extract EMG and we measured peak-to-peak MEP amplitude. Before statistical analysis, we discarded MEPs below resting motor threshold and outside the range of +/- 2 SDs from individual means for each condition (4.71% of data). We normalized the average MEP amplitude for each condition to the rest condition. Two participants (one control and one aphantasic) were removed from the final analysis due to extreme values (outside the range of 2 SDs). The Shapiro-Wilk test was used to check the normality of the data. We did not perform analyses on the timing factor for action reading as we rather isolated the peak of excitability among the four stimulation times for each participant in the action reading condition. This subject-specific peak method accommodates for individual variability in the action semantic processing over time (Dupont et al., 2022a). Finally, to ensure that MEPs were not biased by muscle activation preceding stimulation, for each group, we used Wilcoxon tests to compare the EMGrms before the stimulation artifact between rest and action reading (See Supplementary section). Statistics and data analyses were performed using the Statistica software (Stat Soft, France). The data are presented as mean values (±standard deviation) and the alpha value was set at 0.05.

## Supporting information

Supplementary results

## Author Contributions

Experiment design: WD, CML, FL; Data collection: WD; Statistical analysis: WD, CML, FL; Manuscript preparation: WD, CP, CML, FL.

## Data availability statement

All data from this study are available at https://osf.io/wqm4b/?view_only=96f35d20fefa4db3afd0786dc44f332e

