## Supplementary results for "Mental simulations and action language are impaired in individuals with aphantasia"

### EMGrms:

The absence of muscular pre-activity during action reading was confirmed by a Wilcoxon test revealing no significant difference in EMGrms before TMS artefact between rest and Action reading conditions for Aphantasic ($p=0.534$; $Z=0.620$) and Phantasic individuals ($p=0.097$; $Z=1.654$).

*Table 1: EMGrms activity (mean ±SD) in µV recorded for the first dorsal interosseous before the TMS artifact for each condition (window of 100ms prior the artifact).*

| | **Rest** | **Action reading** |
|---|---|---|
| **Phantasic individuals (Mean ±SD)** | 1.011 ± 0.604 | 1.094 ± 0.510 |
| **Aphantasic individuals (Mean ±SD)** | 1.839 ± 1.422 | 1.639 ± 1.103 |
